# SAINT-Angle: self-attention augmented inception-inside-inception network and transfer learning improve protein backbone torsion angle prediction

**DOI:** 10.1101/2022.12.08.519543

**Authors:** A.K.M. Mehedi Hasan, Ajmain Yasar Ahmed, Sazan Mahbub, M. Saifur Rahman, Md. Shamsuzzoha Bayzid

**Affiliations:** Department of Computer Science and Engineering, Bangladesh University of Engineering and Technology, Dhaka-1205, Bangladesh; Department of Computer Science, University of Maryland, College Park, Maryland 20742, USA

## Abstract

**Motivation:** Protein structure provides insight into how proteins interact with one another as well as their functions in living organisms. Protein backbone torsion angles (*ϕ* and *ψ*) prediction is a key sub-problem in predicting protein structures. However, reliable determination of backbone torsion angles using conventional experimental methods is slow and expensive. Therefore, considerable effort is being put into developing computational methods for predicting backbone angles.

**Results:** We present SAINT-Angle, a highly accurate method for predicting protein backbone torsion angles using a self-attention based deep learning network called SAINT, which was previously developed for the protein secondary structure prediction. We extended and improved the existing SAINT architecture as well as used transfer learning to predict backbone angles. We compared the performance of SAINT-Angle with the state-of-the-art methods through an extensive evaluation study on a collection of benchmark datasets, namely, TEST2016, TEST2018, CAMEO, and CASP. The experimental results suggest that our proposed self-attention based network, together with transfer learning, has achieved notable improvements over the best alternate methods.

**Availability and implementation:** SAINT-Angle is freely available as an open-source project at https://github.com/bayzidlab/SAINT-Angle.

**Supplementary information:** Supplementary material SM.pdf.

## 1 Introduction

Proteins are responsible for various functions in cells and their functions are usually determined by their three-dimensional structures. However, the experimental determination of protein structures using X-ray crystallography, cryogenic electron microscopy (cryo-EM), and nuclear magnetic resonance (NMR) spectroscopy is costly and time- and labour-intensive [1]. Therefore, developing efficient computational approaches for determining protein structures has been gaining increasing attention from the scientific community [2, 3, 4, 5, 6]. The backbone torsion angles (the measurements of the residue-wise torsion [7]) play a critical role in protein structure prediction and investigating protein folding [8]. Therefore, protein structure prediction is often divided into smaller and more doable sub-problems [9] such as backbone torsion angles prediction. As a result, accurate prediction of torsion angles can significantly advance our understanding of the three-dimensional structures of proteins.

Given the growing availability of protein databases and rapid advances in machine learning (ML) methods (especially, the deep learning techniques), application of ML techniques to leverage the available data in accurate prediction of backbone angles has gained significant attention. Earlier ML-based methods used neural network [10, 11], support vector machine (SVM) [11, 12], and hidden Markov model (HMM) [13, 14] to predict discrete states of torsion angles *ϕ* and *ψ*.

Several deep learning-based techniques have recently been developed that can predict backbone torsion angles with a reasonable accuracy. SPIDER2 [15] used iterative neural network to predict the backbone torsion angles, while SPIDER3 [9] leveraged the bidirectional recurrent neural network (BiRNN) [16] to capture the long-range interactions among amino acid residues in a protein molecule. MUFOLD [17] used deep residual inception models [18] to measure the short-range and long-range interactions among different amino acid residues. Similar to SPIDER3, NetSurfP-2.0 [19] used the bidirectional recurrent neural network to capture the long-range interactions. Some studies also emphasized on input feature selection. RaptorX-Angle [20] took advantage of both discrete and continuous representation of the backbone torsion angles and explored the efficacy of different types of features, such as position-specific scoring matrix (PSSM) using PSI-BLAST [21] and position-specific frequency matrix (PSFM) using HHpred [22, 23]. SPOT-1D [24] is an ensemble of nine base models based on the architecture of long short-term memory (LSTM) [25], bidirectional recurrent neural network (BiRNN) [16], and deep residual network (ResNet) [26]. They leveraged the predicted contact-map information produced by SPOT-Contact [27] to improve the performance. OPUS-TASS [6] is another state-of-the-art method, which is an ensemble of 11 base models based on convolutional neural network (CNN) modules [28], bidirectional long short-term memory (BiLSTM) modules [25], and modified Transformer modules [29]. OPUS-TASS is a multi-task learning model [30], maximizing the generalization of neural network, in which the same network was trained for six different prediction tasks. Accuracy of the backbone torsion angles prediction does not rely only on the architecture used in deep learning-based methods but also on the input features extracted from protein sequences. PSSM profiles, HMM profiles [23], physicochemical properties (PP) [31], and amino acid (AA) labels of the residues in proteins are widely used features in predicting protein properties. Recently, the authors of ESIDEN [32] introduced four evolutionary signatures as novel features, namely relative entropy (RE), degree of conservation (DC), position-specific substitution probabilities (PSSP), and Ramachandran basin potential (RBP). ESIDEN is an evolutionary signatures-driven deep neural network developed based on the architecture of long short-term memory (LSTM) and bidirectional long short-term memory (BiLSTM) and achieved notable improvements over other alternate methods.

In this study we present SAINT-Angle, a highly accurate method for protein backbone torsion angle prediction, which is built on our previously proposed architecture of self-attention augmented inception-inside-inception network (SAINT) [33] for protein secondary structure (SS) prediction. We adapted the SAINT architecture for torsion angle prediction and further augmented the basic architecture of SAINT by incorporating the deep residual network [26]. We present a successful utilization of transfer-learning from pretrained transformer-like models by ProtTrans [34] in backbone angle prediction. SAINT-Angle is capable of capturing both short- and long-range interactions among amino acid residues. SAINT was compared with the best alternate methods on a collection of widely used benchmark datasets, namely TEST2016 [27], TEST2018 [24], CAMEO [32], and CASP [32, 33]. SAINT-Angle significantly outperformed other competing methods and achieved the best known *ϕ* and *ψ* prediction accuracy.

## 2 Approach

### 2.1 Feature Representation

SAINT-Angle takes a protein sequence feature vector *X* = {*x*_1_, *x*_2_, …, *x*_*i*_, *x*_*i*+1_….., *x*_*N*_ } as input, where *x*_*i*_ is the vector corresponding to the *i*-th residue of that protein. For each of the residue, SAINT-Angle has 4 regression nodes which predict *sin*(*ϕ*), *cos*(*ϕ*), *sin*(*ψ*), and *cos*(*ψ*) respectively. SAINT-Angle uses three different sets of features which we call 1) *Base features*, 2) *ProtTrans features*, and 3) *Window features*.

The *Base feature* class consists of a feature vector of length 57 for each residue. It contains features from PSSM profiles, HMM profiles, and physicochemical properties (PCP) [31]. We ran PSI-BLAST [21] against the *Uniref90* [35] database with an inclusion threshold of 0.001 and 3 iterations to generate PSSM profiles. We used HHblits [23] using the default parameters against the *uniprot20_2013_03* sequence database to generate the HMM profiles. HHblits also generates seven transition probabilities and three local alignment diversity values, which we used as features as well. Seven physicochemical properties of each amino acid (steric parameters (graph-shape index), polarizability, normalized van der Waals volume, hydrophobicity, isoelectric point, helix probability, and sheet probability) were obtained from [31]. Thus, the dimension of our base feature class for each residue is 57 as this is the concatenation of 20 features from PSSM, 30 features from HMM, and 7 features from physicochemical properties.

The *ProtTrans features*, generated by the pretrained language model for proteins developed by [34], consist of a feature vector of length 1024 for each residue. [34] trained two auto regression language models (Transformer-XL [36] and XLNet [37]) on data containing up to 393 billion amino acids from 2.1 billion protein sequences in a self-supervised manner, considering each residue as a “word” (similar to language modeling in natural language processing [38]). ProtT5-XL-UniRef50 language model has been used in our experiments because it was demonstrated to perform better than other methods on residue-level classification tasks, but other suitable language models from ProtTrans may also be used because our proposed architecture is agnostic about these models. ProtT5-XL-UniRef50 language model generates a sequence of embedding vectors *q* = {*q*_1_, *q*_2_, *q*_3_, …, *q*_*N*_ }, 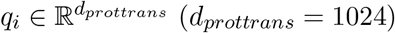 for each residue *X*_*i*_.

*Window features* are generated by windowing the predicted contact information as was done in SPOT-1D and was subsequently used in SAINT. We used SPOT-Contact to generate the contact-maps. We varied the window lengths (the number of preceding or succeeding residues whose pairwise contact information were extracted for a target residue) to generate different dimensional features. We used 4 different window lengths (0, 10, 20, 50) to generate the window features, and denote them by Win0, Win10, Win20, and Win50, respectively.

### 2.2 Architecture of SAINT-Angle

The architecture of SAINT-Angle can be split into three separate discussions: (i) the architecture of SAINT [33], which was proposed for protein secondary structure prediction and our proposed modifications, (ii) the base model architectures of SAINT-Angle that have been applied in an ensemble, and (iii) the overall pipeline of SAINT-Angle.

#### 2.2.1 Architecture of SAINT

Here, we briefly discuss the architecture of SAINT and refer to [33] for details. We have included brief descriptions of the original SAINT architecture to make this paper self-contained and easy to follow. We also discuss the modifications that we have made to the original SAINT architecture to make it suitable for the task of backbone torsion angle prediction. Two of the core components of SAINT are: (i) the self-attention module, and (ii) 2A3I module, which will be discussed in the subsequent subsections.

##### Self-attention module

The self-attention module, as shown in Figure 1(a), that we designed and augmented with the Deep3I network [39] is inspired by the self-attention module developed by [29]. We pass two inputs to our self-attention module: (i) the features from the previous inception module or layer, 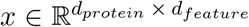, and (ii) position identifiers, 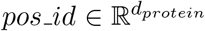, where *d*_*protein*_ is the length of the protein sequence, and *d*_*feature*_ is the length of the feature vector.

**Figure 1:**
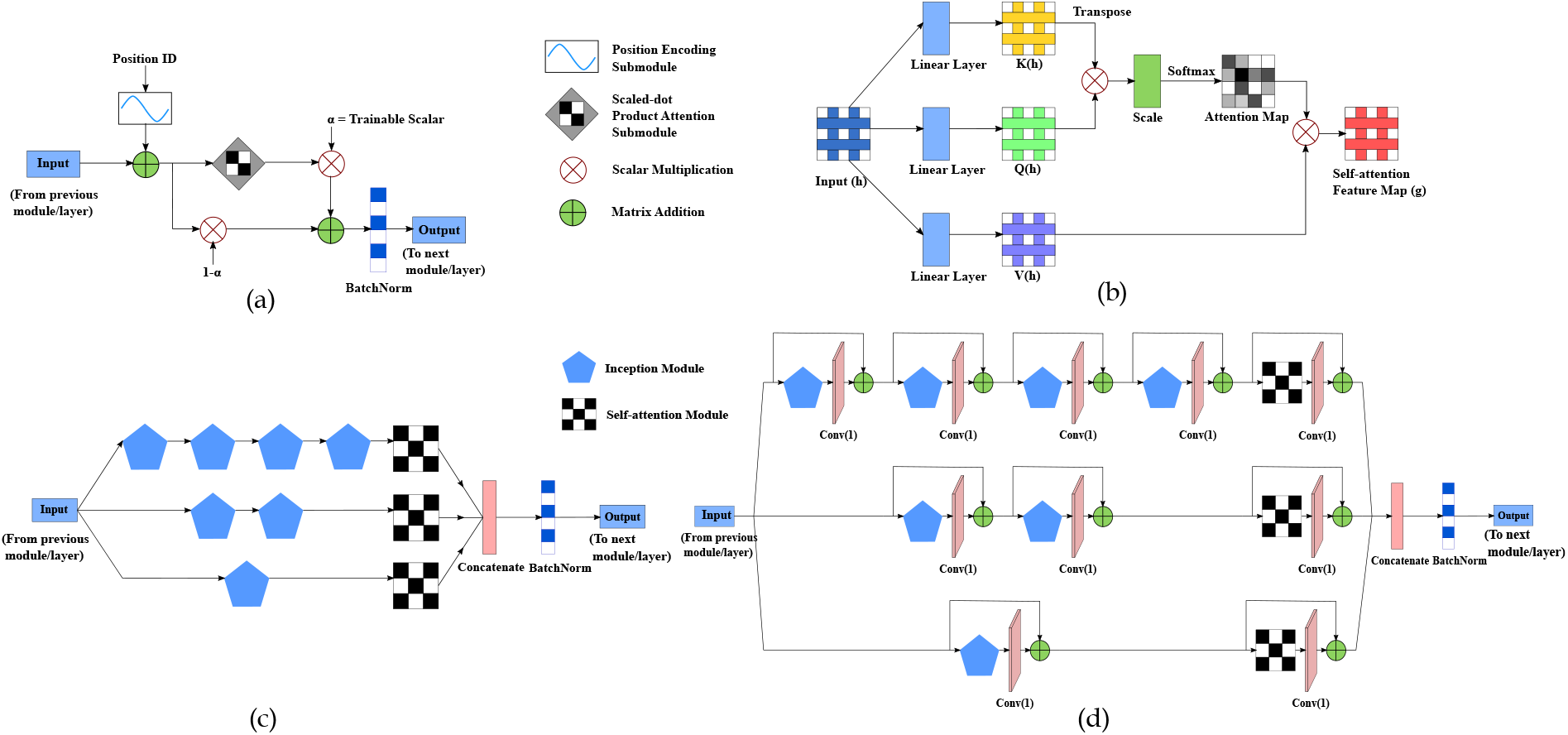
Schematic diagrams of various components of the SAINT-Angle architecture. **(a)** Self-attention module, **(b)** scaled-dot product attention sub-module [29], **(c)** 2A3I module proposed by [33], and **(d)** RES-2A3I module which augments the 2A3I module with residual connections. *Conv*(*x*) denotes a 1-D convolution layer with a kernel of size *x*.

##### Positional encoding sub-module

As the relative or absolute positions of the residues in a protein sequence are important, we need to provide this positional information in our model as shown in Figure 1(a). The *Positional Encoding PosEnc*_*p*_ for a position *p* can be defined as follows [29].

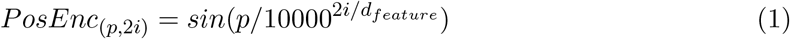

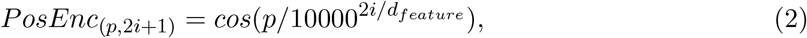

where *i* is the dimension. The above-mentioned function allows the model to learn to attend by relative positions. The inputs *x* is added to the output of positional encoding, resulting in a new representation *h* (Eqn.3). This new representation *h* not only contains the information extracted by the previous layers or modules but also the information about individual positions.

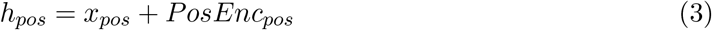

##### Scaled dot-product attention sub-module

We provide the output of the positional encoding sub-module 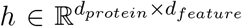 as the input to the *scaled-dot product* attention submodule as shown in Figure 1(b). This input vector is first transformed into three feature spaces *Q, K, V*, representing query, key, and value, respectively. We use three learnable parameter matrices *W*_*Q*_, *W*_*K*_, *W*_*V*_ for this transformation such that *Q*(*h*) = *W*_*Q*_*h, K*(*h*) = *W*_*K*_*h, V* (*h*) = *W*_*V*_ *h*. We then compute the scaled dot-product *s*_*i,j*_ of two vectors *h*_*i*_ and *h*_*j*_ using *Q*(*h*) and *K*(*h*) vectors. This scaled dot-product *s*_*i,j*_ is subsequently used to compute the attention weights 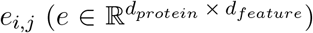, representing how much attention to provide to the vector *I* while synthesizing the vector at position *j*. The output of the scaled dot-product attention sub module *g* is then computed by multiplying the value vector *V* (*h*) with the previously calculated attention weights *e* and subsequently applying batch normalization [40] to reduce the internal covariate shift (Eqn.4). Please see [33] for details.

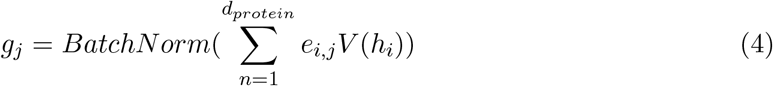

##### 2A3I and RES-2A3I modules

[39] used an assembly of inception modules, which they call 3I (Inception-Inside-Inception) module, in their proposed method MUFOLD-SS to predict protein SS. [33] augmented this with attention modules in order to effectively capture both short- and long-range interactions by placing the self-attention modules (described in Sec. 2.2.1) in each branch of the 3I module as shown in Figure 1(c). This is called the 2A3I (attention augmented inception-inside-inception) module. In this study, we further extended this module by placing residual connections in each of the inception and self-attention modules (Figure 1(d)). Residual connections [26] tackle *vanishing gradient problem* [41] and help make our model more stable. Weight gradients in a neural network are typically very small. During the training of a deep neural network, these small gradients are multiplied by additional small values, resulting in a very small gradient in the earlier layers, and sometimes little or no gradient update at all (as useful gradient information cannot be propagated from the output end of the model back to the layers). This vanishing gradient problem can be addressed by residual connections, producing a more noise stable model with improved learning capacity [42]. We call this residual connection-augmented module the *RES-2A3I* module.

#### 2.2.2 Base Models of SAINT-Angle

We developed the following three architectures that we utilize in an ensemble network to create: (*i*) *Basic* architecture, (*ii*) *ProtTrans* architecture, and (*iii*) *Residual* architecture.

##### Basic architecture

Figure 2(a) shows the schematic diagram of our *Basic* architecture which is identical to the original SAINT architecture proposed for protein SS prediction [33]. It starts with two consecutive 2A3I modules followed by a self-attention module. This self-attention module supplements the amount of non-local interactions that have been captured by previous two 2A3I modules. Next, we have an 1D convolutional layer with window size 11. The output of the convolutional layer is passed through another self-attention module followed by two dense layers, with yet another self-attention module placed in between these two dense layers. This self-attention module helps in understanding how the residues align and interact, making it easier to comprehend the behavior of the model. The final dense layer has four regression nodes that infer *sin*(*ϕ*), *cos*(*ϕ*), *sin*(*ψ*), and *cos*(*ψ*).

**Figure 2:**
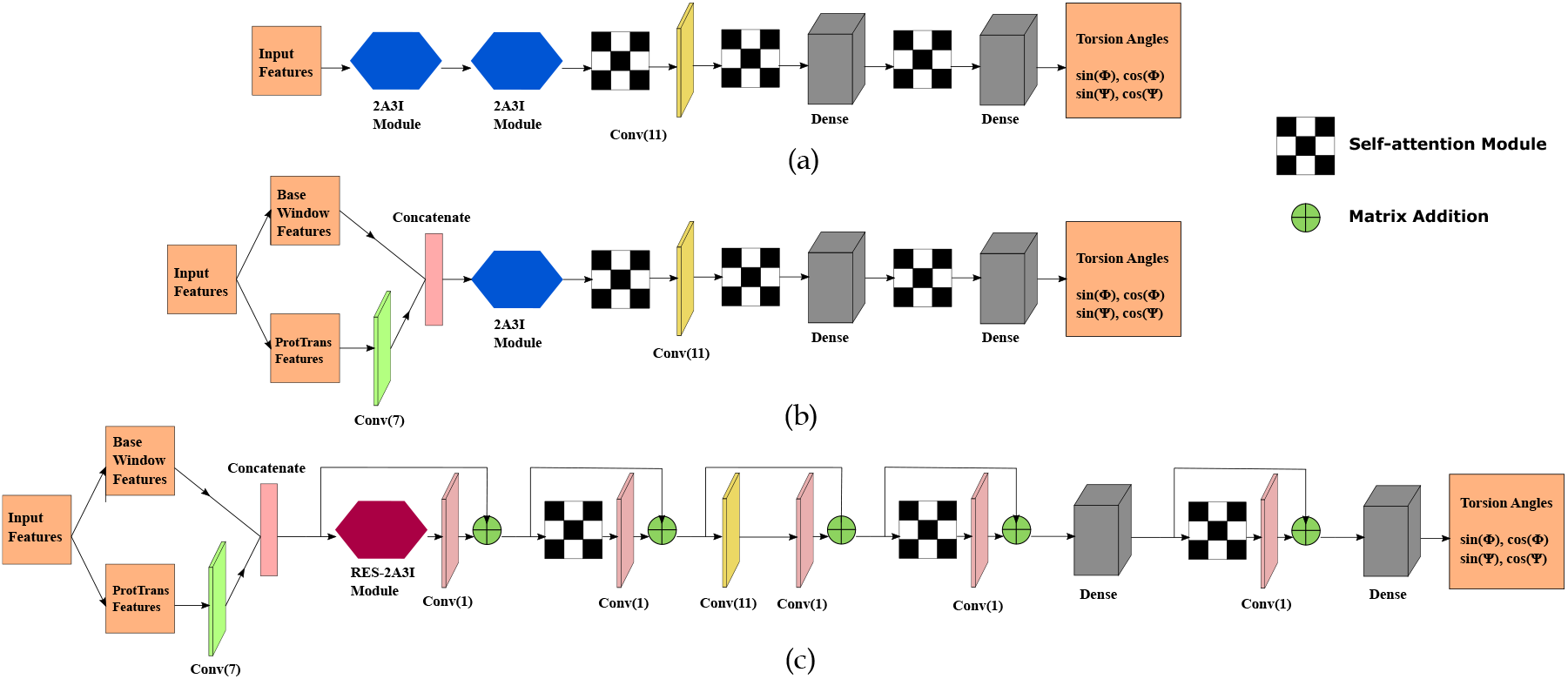
Schematic diagrams of the architectures of the base models used in SAINT-Angle. **(a)** The basic architecture which was proposed by [33], **(b)** the ProtTrans architecture, and **(c)** the residual architecture which augments the ProtTrans architecture with residual connections.

##### ProtTrans architecture

We developed the ProtTrans architecture (Figure 2 (b)) to effectively use the ProtTrans features by treating them differently from the base and window features. We pass the ProtTrans features to a 1D convolution layer with window size 7. This convolutional layer acts as a local feature extractor, capturing local interactions between residues and reducing the dimension of the ProtTrans features from a 1024-dimensional feature vector to a 300-dimensional feature vector, allowing the model to filter out less important information. It also aids in avoiding over-fitting and reducing the number of trainable parameters. The output of this 1D convolution layer is then concatenated with the base and window features. The concatenated vector is then passed through a single 2A3I module. Note that, unlike the basic SAINT architecture, we have only one 2A31 module as we observed that two 2A31 modules do not provide notable advantage in this architecture but increases the training time. The rest of the architecture is similar to the basic SAINT architecture.

##### Residual architecture

The *Residual* architecture (Figure 2(c)) is similar to the *ProtTrans* architecture except for two differences: (i) we have added residual connections [26] between different components as shown in Figure 2 (c), and (ii) we have used the RES-2A3I module instead of the 2A3I module. Residual connections enable the deeper layers to use the features extracted from the earlier layers. Usually, the deeper level layers use features that are highly convoluted and lower in resolution. Residual connections help the deeper layers leverage the low-level and high dimensional features. It also helps to make the model stable.

#### 2.2.3 Overview of SAINT-Angle

SAINT-Angle is an ensemble network of eight models with different combinations of architectures (discussed in Sec. 2.2.2) and features (discussed in Sec. 2.1) – resulting in a set of diverse learning paths leveraging different types of features. Table 1 shows the architectures and features used in these eight models. Details of the ensemble and the individual models therein are presented in Supplementary Material. We used the same training and validation sets that were used by SPOT-1D to train these models and tune necessary hyperparameters. Each model was trained using Adam optimizer [43] with an initial learning rate of 0.0001 which was subsequently reduced by half when the accuracy of the validation set did not improve for five consecutive epochs.

**Table 1:** Eight models used in SAINT-Angle. We show the architectures and features used in these eight models.

| Model | Architecture | Feature set |
| --- | --- | --- |
| 1 | Basic | Base+Win0 |
| 2 | Basic | Base+Win10 |
| 3 | ProtTrans | Base+ProtTrans+Win0 |
| 4 | ProtTrans | Base+ProtTrans+Win10 |
| 5 | ProtTrans | Base+ProtTrans+Win20 |
| 6 | ProtTrans | Base+ProtTrans+Win50 |
| 7 | Residual | Base+ProtTrans+Win0 |
| 8 | Residual | Base+ProtTrans+Win10 |

## 3 Results and Discussion

We performed an extensive evaluation study, comparing SAINT-Angle with the recent state-of-the-art methods on a collection of widely used benchmark dataset.

### 3.1 Dataset

#### 3.1.1 Training and validation dataset

The SPOT-1D dataset [24, 27] was used for training and validation of SAINT-Angle. These proteins were culled from the PISCES server [44] on February 2017 with resolution *<*2.5 Å, *R* − *free <*1, and a sequence identity cutoff of 25% according to BlastClust [21]. The proteins consisting of over 700 amino acid residues were removed by the authors of SPOT-1D to fit in the SPOT-Contact [27] pipeline. As a result, 10,029 and 983 proteins remained in the training and validation sets respectively.

#### 3.1.2 Test dataset

We assessed the performance of SAINT-Angle and other competing methods on a collection of widely used test sets, which are briefly described in Sec. 3 in the Supplementary Material.

### 3.2 Performance evaluation

We compared our proposed SAINT-Angle with several state-of-the-art methods: OPUS-TASS, SPOT-1D, NetSurfP-2.0, MUFOLD, SPIDER3, and RaptorX-Angle. We also compared SAINT-Angle with a recent and highly accurate method ESIDEN [32]. We evaluated the performance of backbone torsion angles prediction methods using mean absolute error (MAE) which is the measure of average absolute difference between native values (*T*) and predicted values (*P*) over all amino acid residues in a protein. The minimum value between |*T* − *P* | and 360^◦^ − |*T* − *P* | was taken in order to reduce the periodicity of an angle [32] as given in Eqn.5. Here *N* is the number of proteins, *L*_*i*_ is the total number of amino acid residues in the *i*th protein. *T*_*ij*_ and *P*_*ij*_ are the values of native and predicted angles of the *j*-th amino acid residue in the *i*-th protein respectively. We performed Wilcoxon signed-rank test [45] (with *α* = 0.05) to measure the statistical significance of the differences between two methods.

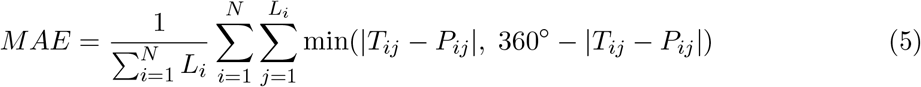

### 3.3 Results on benchmark datasets

The comparison of SAINT-Angle with other state-of-the-art methods on TEST2016 and TEST2018 is shown in Table 2. Experimental results show that SAINT-Angle outperforms all other methods on both TEST2016 and TEST2018 datasets. Especially, the improvement in MAE(*ψ*) is substantial – almost two degrees over the second best method OPUS-TASS, and more than 2-6 degrees compared to other methods. Notably, even the individual eight base models used in SAINT-Angle achieves comparable or better performance than most other methods (see Table S4 in Supplementary Material). Among these individual base models, the improvements of the models with *ProtTrans* architecture and ProtTrans features (Models 3, 4, 5, and 6 in Table 1) over the *Basic* architecture with base features (Models 1 and 2) are notable – indicating the successful utilization of ProtTrans-based transfer learning. Statistical tests suggests that these improvements of SAINT-Angle over other methods are statistically significant (*p* − *value* ≪ 0.05). RaptorX-Angle performed poorly compared to other methods on both these datasets. OPUS-TASS and SPOT-1D produced similar results, with OPUS-TASS obtaining marginally better results than SPOT-1D. Both OPUS-TASS and SPOT-1D obtained notably better results than NetSurfP, MUFOLD and SPIDER3.

**Table 2:** Performance (in terms of MAE(*ϕ*) and MAE(*ψ*)) of SAINT-Angle and other state-of-the-art methods on TEST2016 and TEST2018. The best and the second best results are shown in bold and italic, respectively. Values which were not reported by the corresponding source are indicated by “-”.

| Method | TEST2016 |  | TEST2018 |  |
| --- | --- | --- | --- | --- |
| | $MAE(\phi)$ | $MAE(\psi)$ | $MAE(\phi)$ | $MAE(\psi)$ |
| RaptorX-Angle<sup>a</sup> | 18.08 | 26.68 | 21.01 | 35.95 |
| SPIDER3<sup>a</sup> | 17.88 | 26.66 | 18.38 | 28.10 |
| MUFOLD<sup>a</sup> | - | - | 17.78 | 27.24 |
| NetSurfP-2.0<sup>a</sup> | - | - | 17.90 | 26.63 |
| SPOT-1D<sup>a</sup> | 16.27 | 23.26 | 16.89 | 24.87 |
| OPUS-TASS<sup>b</sup> | *15.78* | *22.46* | *16.40* | *24.06* |
| SAINT-Angle | **15.45** | **20.77** | **16.25** | **22.37** |

<sup>a</sup>Results reported by SPOT-1D [24].
<sup>b</sup>Results reported by OPUS-TASS [6].

The performance of SAINT-Angle and other competing methods on three CASP datasets (CASP12, CASP13, and CASP-FM) is shown in Table 3. SAINT-Angle consistently outperformed other methods on these datasets, with only one exception on CASP-FM where OPUS-TASS obtained a better MAE(*ϕ*) than SAINT-Angle albeit the difference is very small (0.01 degree). Similar to TEST2016 and TEST2018 datasets, the improvements of SAINT-Angle over other methods in MAE(*ψ*) are more substantial than those in MAE(*ϕ*). The improvements of SAINT-Angle over other methods are statistically significant (*p* − *value* ≪ 0.05).

**Table 3:** Performance (in terms of MAE(*ϕ*) and MAE(*ψ*)) of SAINT-Angle and other state-of-the-art methods on on three CASP datasets. The best and the second best results are shown in bold and italic, respectively.

| Method | CASP12 (55) |  | CASP13 (32) |  | CASP-FM (56) |  |
| --- | --- | --- | --- | --- | --- | --- |
| | $\text{MAE}(\phi)$ | $\text{MAE}(\psi)$ | $\text{MAE}(\phi)$ | $\text{MAE}(\psi)$ | $\text{MAE}(\phi)$ | $\text{MAE}(\psi)$ |
| SPOT-1D<sup>a</sup> | 18.44 | 26.90 | 18.48 | 26.73 | 19.39 | 30.10 |
| OPUS-TASS<sup>a</sup> | *18.08* | *25.98* | *17.89* | *25.93* | **18.85** | *28.00* |
| SAINT-Angle | **17.91** | **24.76** | **17.55** | **23.96** | *18.86* | **27.65** |

<sup>a</sup>Results reported in OPUS-TASS.

### 3.4 Comparison of SAINT-Angle with ESIDEN

ESIDEN [32] – a recent, highly accurate recurrent neural network based method – introduced and leveraged four evolutionary signatures as novel features, namely relative entropy (RE), degree of conservation (DC), position-specific substitution probabilities (PSSP), and Ramachandran basin potential (RBP). They showed that these novel features, along with classical features such as PSSM, physicochemical properties (PP), and amino acid (AA), result in significant improvements in protein torsion angle prediction. As ESIDEN is not an ensemble-based network, in order to make a fair comparison with ESIDEN and to further assess the efficacy of the novel evolutionary features, we trained our *basic architecture* (discussed in Sec. 2.2.2) using the features used by ESIDEN and evaluated its performance on a collection of datasets compiled and analyzed by the authors of ESIDEN. This will enable us to assess the performance of the basic SAINT architecture using the features used by ESIDEN (i.e., without the ProtTrans-based transfer learning and the ensemble network). We call this basic architecture with ESIDEN features *SAINT-Angle-Single*. We obtained the evolutionary features for the SPOT-1D training dataset and a collection of test datasets from the authors of ESIDEN. We also trained our ensemble network using the base features along with the novel ESIDEN features (i.e, 20 types of amino acids (AA) and four evolutionary features DC, RE, PSSP, and RBP). The evolutionary features of ESIDEN were shown to be reasonably powerful [32], which has been further supported by our experimental results as well (discussed later in this section). On the other hand, the window features are difficult to compute [6]. Therefore, in these experiments, we did not use the window features to keep the dimension of the feature vector manageable as well as to best take advantage of the evolutionary features used in ESIDEN. Thus, after removing the window features when ESIDEN features are available, we had three models (out of eight models listed in Table 1) to use for the ensemble network (see Sec. 2.1 and Table S5 in Supplementary Material). In order to distinguish this ensemble of three base models using the ESIDEN features from the ensemble of eight models, we call this SAINT-Angle^∗^.

The comparison of SAINT-Angle-Single, SAINT-Angle^∗^ (ensemble of three models using the ESIDEN features), and SAINT-Angle (ensemble of eight models without ESIDEN features) with ESIDEN on TEST2016 and TEST2018 datasets is shown in Table 4. ESIDEN is notably better than the SPOT-1D, OPUS-TASS as well as SAINT-Angle, especially for predicting the *ψ* angle (around 4^◦^ improvement in MAE(*ψ*)). Note however that SAINT-Angle, unlike ESIDEN, does not use the four evolutionary features. Interestingly, SAINT-Angle-Single, which leverages ESIDEN features, is remarkably better than ESIDEN (∼ 2^◦^ improvement in MAE(*ψ*)). This shows the superiority of our SAINT architecture over the ESIDEN architecture. The performance of SAINT-Angle-Single and SAINT-Angle^∗^ is mixed on these two datasets. SAINT-Angle^∗^ is better than SAINT-Angle-Single on TEST2016 dataset whereas SAINT-Angle-Single is better than SAINT-Angle^∗^ on TEST2018. Remarkably, both of them achieved substantial improvements over ESIDEN. Moreover, both SAINT-Angle-Single and SAINT-Angle^∗^ outperformed SAINT-Angle – showing the power of the evolutionary features proposed by ESIDEN. The improvements of SAINT-Angle-Single and SAINT-Angle^∗^ over ESIDEN and SAINT-Angle are statistically significant (*p* − *value* ≪ 0.05).

**Table 4:** Performance of SAINT-Angle-Single (basic SAINT architecture with base and ESIDEN features), SAINT-Angle^∗^ (ensemble of three models using the ESIDEN features), and SAINT-Angle (ensemble of eight models without ESIDEN features), and ESIDEN on TEST2016 and TEST2018.

| Method | TEST2016 |  | TEST2018 |  |
| --- | --- | --- | --- | --- |
| | $\text{MAE}(\phi)$ | $\text{MAE}(\psi)$ | $\text{MAE}(\phi)$ | $\text{MAE}(\psi)$ |
| SPOT-1D | 16.27 | 23.26 | 16.89 | 24.87 |
| OPUS-TASS | 15.78 | 22.46 | 16.40 | 24.06 |
| ESIDEN<sup>a</sup> | 15.48 | 19.25 | 16.00 | 20.28 |
| SAINT-Angle | 15.45 | 20.77 | 16.25 | 22.37 |
| SAINT-Angle-Single | *14.39* | *17.03* | **14.89** | **17.94** |
| SAINT-Angle* | **14.18** | **16.59** | *15.48* | *18.77* |

<sup>a</sup>Results reported by ESIDEN [32].

We further assessed the performance of SAINT-Angle-Single in comparison with ESIDEN and other methods on five other benchmark datasets that were compiled and used by the authors of ESIDEN, namely CAMEO109 and four CASP datasets (CASP11, CASP12, CASP13, CASP14). Note that these CASP datasets (analyzed in ESIDEN [32]) are different from the CASP datasets in Table 3(which was used by SAINT). Results on the CAMEO109 dataset are shown in Table 5. ESIDEN is better than other existing methods in terms of MAE(*ψ*) (∼ 1^◦^ improvement), but SPOT-1D obtained slightly better MAE(*ϕ*) than ESIDEN. SAINT-Angle-Single outperformed ESIDEN and other methods in terms of both MAE(*ϕ*) and MAE(*ψ*). Especially, it obtained around two degrees of improvement over ESIDEN in MAE(*ψ*).

**Table 5:** Performance of SAINT-Angle-Single and other state-of-the-art methods on CAMEO109.

| Method | CAMEO109 |  |
| --- | --- | --- |
| | MAE($\phi$) | MAE($\psi$) |
| RaptorX-Angle<sup>a</sup> | 19.57 | 33.96 |
| SPIDER3<sup>a</sup> | 17.89 | 28.32 |
| SPOT-1D<sup>a</sup> | *16.49* | 25.17 |
| ESIDEN<sup>a</sup> | 16.57 | *24.25* |
| SAINT-Angle-Single | **16.40** | **22.77** |

<sup>a</sup>Results reported by ESIDEN.

Results on four CASP datasets are shown in Table 6. SAINT-Angle-Single and ESIDEN are significantly better than other existing methods, especially for *ψ* where ESIDEN and SAINT-Angle achieved more than ∼ 10^◦^ improvements over other methods. Remarkably, SAINT-Angle-Single outperformed all other methods (including ESIDEN) across all the datasets in terms of both MAE(*ϕ*) and MAE(*ψ*), with only one exception where ESIDEN obtained a better MAE(*ϕ*) than SAINT-Angle-Single on the CASP11 dataset.

**Table 6:** Performance of SAINT-Angle-Single and other state-of-the-art methods on four CASP datasets.

| Method | CASP11 (27) |  | CASP12 (11) |  | CASP13 (13) |  | CASP14 (8) |  |
| --- | --- | --- | --- | --- | --- | --- | --- | --- |
| | MAE($\phi$) | MAE($\psi$) | MAE($\phi$) | MAE($\psi$) | MAE($\phi$) | MAE($\psi$) | MAE($\phi$) | MAE($\psi$) |
| RaptorX-Angle<sup>a</sup> | 20.33 | 40.05 | 21.71 | 38.22 | 22.73 | 41.18 | 24.95 | 48.06 |
| SPIDER3<sup>a</sup> | 19.19 | 34.63 | 21.14 | 34.92 | *22.48* | 38.46 | 23.41 | 38.79 |
| SPOT-1D<sup>a</sup> | 18.54 | 25.77 | 20.21 | 31.71 | 22.60 | 34.28 | 23.42 | 33.44 |
| ESIDEN<sup>a</sup> | **17.25** | *23.30* | *19.94* | *28.86* | **22.15** | *32.40* | *23.01* | *29.96* |
| SAINT-Angle-Single | *17.40* | **22.52** | **19.58** | **27.11** | **22.15** | **31.46** | **22.67** | **29.83** |

*Note:* The numbers of proteins in these datasets are shown in parentheses.
<sup>a</sup>Results reported by ESIDEN.

### 3.5 Analysis of the predicted angles

We further investigated the predicted protein backbone torsion angles from SAINT-Angle and other contemporary methods to obtain better insights on the performances of various methods.

#### 3.5.1 Impact of long-range interactions

We investigated the effect of long-range interactions among amino acid residues in protein torsion angle prediction. Two residues at sequence position *i* and *j* are considered to have non-local contact or interaction if they are at least twenty residues apart (|*i* − *j*|≥ 20), but *<*8 Å away in terms of atomic distance between their alpha carbon (*C*_*α*_) atoms [9]. We computed the average number of non-local interactions per residue for each of the 1213 target proteins in the TEST2016 dataset and sorted the proteins in an ascending order of their average number of non-local interactions per residue. Next, we put them in six equal-sized bins (*b*_1_, *b*_2_, …, *b*_6_) where the first bin contained the proteins with the lowest level of non-local interactions (0-0.61 non-local contacts per residue) and the sixth bin contains the proteins with the highest level of non-local interactions (1.64-2.70 non-local contacts per residue). Figure 3 shows the MAE(*ϕ*) and MAE(*ϕ*) for the best performing methods for these six bins.

**Figure 3:**
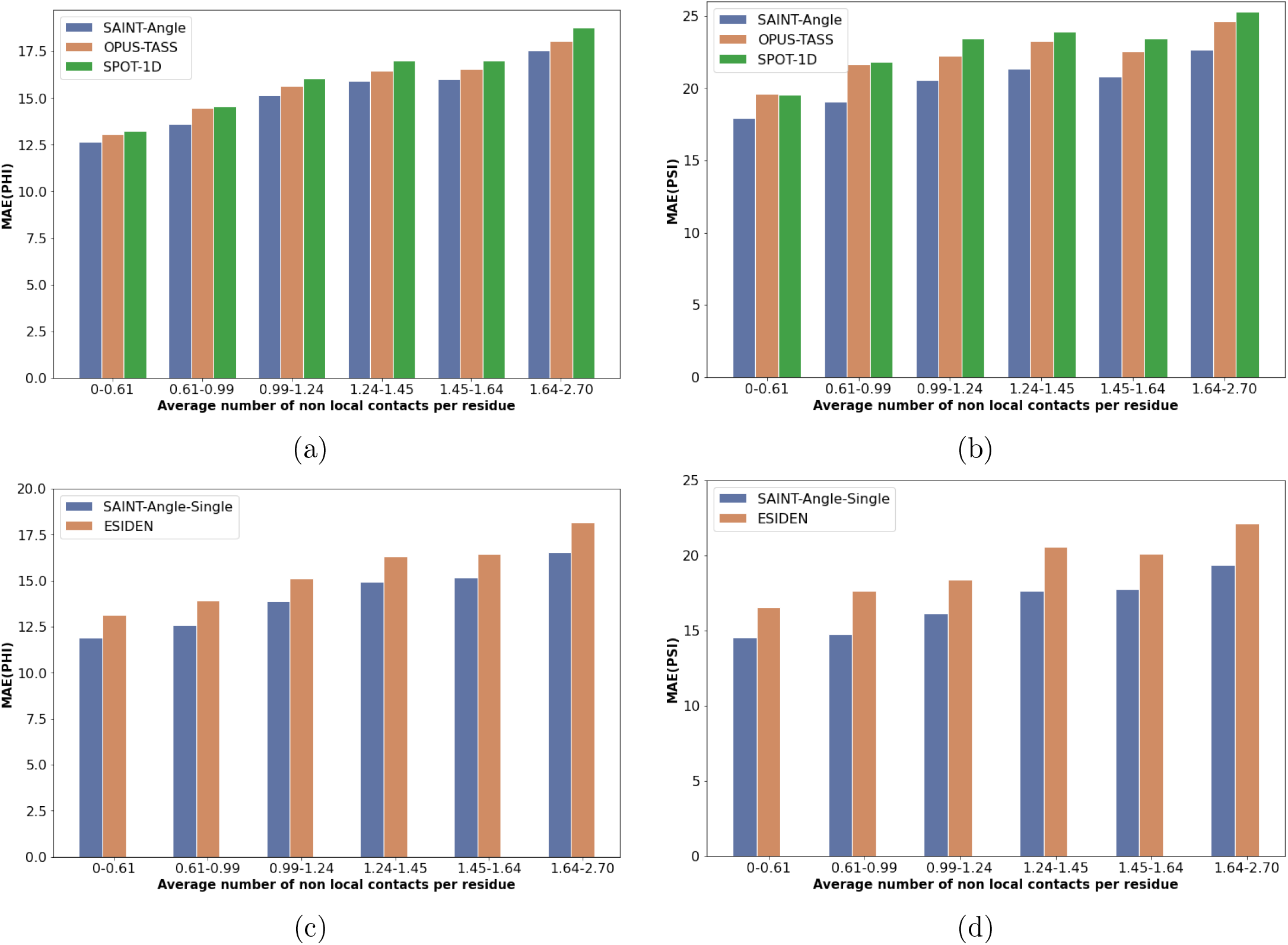
MAE(*ϕ*) and MAE(*ϕ*) of SAINT-Angle and the best alternative methods under various levels of non-local interactions. We show the results on the TEST2016 test set using six bins of proteins. **(a)** MAE(*ϕ*) of SAINT-Angle, OPUS-TASS, and SPOT-1D under various levels of non-local contacts. **(b)** MAE(*ψ*) of SAINT-Angle, OPUS-TASS, and SPOT-1D under various levels of non-local contacts. **(c)-(d)** MAE(*ϕ*) and MAE(*ψ*) of SAINT-Angle-Single and ESIDEN under various levels of non-local contacts.

These results show that – as expected – the performance of SAINT-Angle and other methods degrades as we increase the number of non-local contacts. However, SAINT-Angle is consistently and significantly (*p* − *value* ≪ 0.05) better than the best alternate methods across all levels of non-local interactions. Moreover, the improvements of SAINT-Angle (or SAINT-Angle-Single) over other methods tend to gradually increase with increasing levels of non-local interactions from *b*_1_ to *b*_6_ (with a few exceptions), especially for the torsion angle *ψ*. Similarly, there is no notable difference between SPOT-1D and OPUS-TASS on *b*_1_, whereas there are notable differences on *b*_6_. These results indicate that long-range interactions have an impact on torsion angle prediction, and that capturing non-local interactions by self-attention modules is one of the contributing factors in the improvement of SAINT-Angle.

#### 3.5.2 Impact of 8-class (Q8) secondary structure states

We analyzed the performance of various methods on the 1213 target proteins in the TEST2016 dataset across eight types of secondary structure states, namely *β*-bridge (B), coil (C), *β*-strand (E), 3_10_-helix (G), *α*-helix (H), *π*-helix (I), bend (S), and *β*-turn (T). Figure S1 in Supplementary Material shows the average MAE(*ϕ*) and MAE(*ψ*) against each of the Q8 labels for various methods. These results suggest that H (*α*-helix), G (3_10_-helix), E (*β*-strand), and I (*π*-helix) regions usually have lower prediction errors whereas the non-ordinary states [46], such as S (bend), C (coil), B (*β*-bridge) and T (*β*-turn) regions generally have higher prediction errors. Another notable observation is that SAINT-Angle consistently obtained superior performance across all Q8 labels compared to its competing methods, except that OPUS-TASS and ESIDEN obtained marginally better average MAE(*ψ*) values than SAINT-Angle on I states (see Fig. S1(b) and (d)). Note that the I (*π*-helix) secondary state is extremely rare, appearing in only about 15% of all known protein structures, and is difficult to predict [47]. Notably, while the performances of different methods on the easy regions (e.g., H regions) are comparable, SAINT-Angle is notably better than other methods on regions where angle prediction is relatively hard (e.g., S and T regions).

## 4 Conclusions

We have presented SAINT-Angle, a highly accurate method for protein backbone angle (*ϕ* and *ψ*) prediction. We have augmented the basic SAINT architecture for effective angle prediction and showed a successful utilization of transfer learning from pre-trained transformer-like language models. SAINT-Angle was assessed for its performance against the state-of-the-art backbone angle prediction methods on a collection of widely used benchmark datasets. Experimental results suggest that SAINT-Angle consistently improved upon the best existing methods.

The self-attention module in the SAINT architecture was particularly aimed for effectively capturing long-range interactions, and our systematic analyses of the performance of different methods under various model conditions with varying levels of long-range interactions indicate that SAINT-Angle can better handle complex models conditions with high level of long-range interactions. The improvement of SAINT-Angle over other methods in *ψ* prediction, which is typically harder to predict than *ϕ*, is noteworthy as it achieved more than 2-6 degree less MAE(*ψ*) than other methods on benchmark datasets. To the best of our knowledge, we – for the first time – leveraged ProtTrans features for angle prediction. The ProtTrans architecture (discussed in Sec. 2.2.2) alone (i.e., without the ensemble network) performs better than the existing best methods like OPUS-TASS and SPOT-1D – showing the positive impact of transfer learning in protein attribute prediction. We also analyzed the novel evolutionary features proposed in [32]. Our analyses with the evolutionary features reconfirms the effectiveness of the evolutionary features in protein angle prediction which was first demonstrated by [32]. Our results also suggest that the architecture of SAINT-Angle is *feature-robust*, as it performs well with different types of features (e.g., ProtTrans features, evolutionary features proposed by [32]) and consistently outperforms other competing methods on varying feature sets. Given this demonstrated performance improvement on various benchmark datasets and under challenging model conditions, we believe SAINT-Angle advances the state-of-the-art in this domain, and will be considered as a useful tool for predicting the backbone torsion angles.

This study can be extended in several directions. As an immediate future direction, we plan to train a multi-task learning model by leveraging the original SAINT and the proposed SAINT-Angle architectures, which would simultaneously predict protein secondary structures and torsion angles. Although SAINT-Angle appears to be feature-robust, follow up studies need to investigate various features and select a small set of simple features that are sufficient for SAINT-Angle to predict backbone angles with a reasonable accuracy. Another research direction is to extend and utilize the proposed model to predict other protein attributes.

## Supplementary Materials

### 1 Overview

These supplementary materials present additional details about the base models used in our ensemble network (Sec. 2), the datasets used to validate and test our models (Sec. 3), and some additional results (Sec. 4).

### 2 Base Models in Ensemble Network

We initially trained 24 different models with different feature sets and model architectures. The same training and validation sets that were used by SPOT-1D [1] were used to train these models and tune necessary hyperparameters. Different combinations of model architectures and feature sets were explored. In addition to the base, ProtTrans, and window features (as discussed in Sec. 2.1 in the main paper), we made an attempt to use the native 8-class (Q8) secondary structure (SS) of the residues as a feature, which resulted in a significant improvement in backbone angle prediction (results not shown). However, native structure is not available for all proteins, especially for the newly discovered ones. So, we used SAINT [2] to produce predicted probability distribution of the Q8 states of the residues in a given protein. We call this predicted Q8 states the *predicted features*. Thus, we used 16 different feature sets as shown in Table S1. As we will show later, unlike the true SS, using the predicted features did not result in notable improvements in backbone angle prediction.

We used the basic architecture on feature sets *fs*1 − *fs*8, thereby producing eight models. On features sets *fs*9 − *fs*16, we applied both the ProtTrans and residual architectures, resulting in 16 different models. Thus, in total, we trained 24 models. Hence, we obtained 24 models in total. We evaluated the performance of each of these models on the validation set (see Table S2). Besides, we conducted an inclusion-exclusion study to select a set of models for an ensemble network. We did not observe any advantage of using the predicted features. Finally, we selected 8 models, as shown in Table S3 to form the ensemble model which we call SAINT-Angle. Table S4 shows the comparison of these based models with existing best performing methods on TEST2016 and TEST2018 benchmark datasets.

**Table S1:** Different feature sets and their acronyms.

| Feature set | Notation | Feature set | Notation |
| --- | --- | --- | --- |
| Base+Win0 | fs1 | Base+ProtTrans+Win0 | fs9 |
| Base+Win10 | fs2 | Base+ProtTrans+Win10 | fs10 |
| Base+Win20 | fs3 | Base+ProtTrans+Win20 | fs11 |
| Base+Win50 | fs4 | Base+ProtTrans+Win50 | fs12 |
| Base+Win0+Predicted | fs5 | Base+ProtTrans+Win0+Predicted | fs13 |
| Base+Win10+Predicted | fs6 | Base+ProtTrans+Win10+Predicted | fs14 |
| Base+Win20+Predicted | fs7 | Base+ProtTrans+Win20+Predicted | fs15 |
| Base+Win50+Predicted | fs8 | Base+ProtTrans+Win50+Predicted | fs16 |

**Table S2:** Performance of the 24 individual models on the validation set.

| Feature | Basic architecture |  | Feature | ProtTrans Architecture |  | Residual Architecture |  |
| --- | --- | --- | --- | --- | --- | --- | --- |
| | MAE($\phi$) | MAE($\psi$) | | MAE($\phi$) | MAE($\psi$) | MAE($\phi$) | MAE($\psi$) |
| fs1 | 16.77 | 23.69 | fs9 | 15.74 | 21.17 | 15.73 | 21.23 |
| fs2 | 16.38 | 22.74 | fs10 | 15.63 | 20.67 | 15.56 | 20.67 |
| fs3 | 16.35 | 22.79 | fs11 | 15.64 | 20.75 | 15.61 | 20.80 |
| fs4 | 16.41 | 22.80 | fs12 | 15.69 | 20.74 | 15.75 | 20.94 |
| fs5 | 16.43 | 22.85 | fs13 | 15.70 | 20.90 | 15.63 | 20.83 |
| fs6 | 16.40 | 22.89 | fs14 | 15.68 | 21.05 | 15.61 | 20.95 |
| fs7 | 16.44 | 22.85 | fs15 | 15.72 | 20.97 | 15.61 | 21.05 |
| fs8 | 16.43 | 22.86 | fs16 | 15.71 | 21.09 | 15.68 | 21.04 |

**Table S3:** Performance of the selected eight base models and the corresponding ensemble network on the validation set.

| Model | Feature set | Architecture | validation error |  |
| --- | --- | --- | --- | --- |
| | | | MAE($\phi$) | MAE($\psi$) |
| Model 0 | fs1 | Basic | 16.77 | 23.69 |
| Model 1 | fs2 | Basic | 16.38 | 22.74 |
| Model 2 | fs9 | ProtTrans | 15.74 | 21.17 |
| Model 3 | fs10 | ProtTrans | 15.63 | 20.67 |
| Model 4 | fs11 | ProtTrans | 15.64 | 20.75 |
| Model 5 | fs12 | ProtTrans | 15.69 | 20.74 |
| Model 6 | fs9 | Residual | 15.73 | 21.23 |
| Model 7 | fs10 | Residual | 15.56 | 20.67 |
| Ensemble |  |  | 15.39 | 20.14 |

**Table S4:** Performance (in terms of MAE(*ϕ*) and MAE(*ψ*)) of the individual base models used in SAINT-Angle and the best competing methods on TEST2016 and TEST2018. The best and the second best results are shown in bold and italic, respectively.

| Method | TEST2016 |  | TEST2018 |  |
| --- | --- | --- | --- | --- |
| | MAE($\phi$) | MAE($\psi$) | MAE($\phi$) | MAE($\psi$) |
| Model 0 | 16.80 | 24.38 | 17.37 | 25.93 |
| Model 1 | 16.51 | 23.70 | 16.99 | 25.20 |
| Model 2 | 15.95 | 21.88 | 16.18 | 23.04 |
| Model 3 | *15.76* | **21.45** | *16.16* | **22.71** |
| Model 4 | 15.79 | *21.50* | **16.13** | *22.76* |
| Model 5 | 15.84 | 21.54 | 16.21 | 22.78 |
| Model 6 | 15.91 | 22.02 | 16.58 | 23.47 |
| Model 7 | **15.75** | 21.57 | 16.58 | 23.06 |
| SPOT-1D<sup>a</sup> | 16.27 | 23.26 | 16.89 | 24.87 |
| OPUS-TASS<sup>b</sup> | 15.78 | 22.46 | 16.40 | 24.06 |

<sup>a</sup>Results reported by SPOT-1D [1].
<sup>b</sup>Results reported by OPUS-TASS [3].

#### 2.1 Ensemble using the features proposed by ESIDEN

In order to compare with ESIDEN [4] and to take advantage of the novel features proposed by ESIDEN, we removed the window features and used the ESIDEN features instead – resulting in three models as shown in Table S5. So, in this case, the feature set contains 3 basic features including 20 types of amino acids (AA), 7 physicochemical properties (PCP), and position-specific scoring matrix (PSSM) along with 4 novel evolutionary features proposed by ESIDEN, namely degree of conservation (DC), relative entropy (RE), position-specific substitution probabilities (PSSP), and Ramachandran basin potential (RBP). We call this variant, which is an ensemble of the three models as shown in Table S5, SAINT-Angle ^∗^.

**Table S5:** Three models used in SAINT-Angle^∗^ when the ESIDEN features are available. We show the architectures and features used in these three models.

| Model | Architecture | Feature set |
| --- | --- | --- |
| 0 | Basic | Base (PSSM, HMM, PCP) + ESIDEN (AA, DC, RE, PSSP, RBP) |
| 1 | ProtTrans | Base+ESIDEN+ProtTrans |
| 2 | Residual | Base+ESIDEN+ProtTrans |

### 3 Test Datasets

**TEST2016** TEST2016 was compiled by [5], containing 1213 proteins that were deposited on PDB [6] between June 2015 and February 2017 with similar parameter settings as the training and validation sets. None of the proteins contains more than 700 amino acid residues. The dataset has *<*25% sequence similarity with the training and validation sets according to BlastClust.

**TEST2018** TEST2018 dataset, compiled by [5], contains 250 high-quality, non-redundant proteins that were deposited in PDB between January 2018 and July 2018 with the same parameter settings and filtering constraints as the TEST2016 dataset.

**CAMEO109** CAMEO109 dataset contains 109 proteins that were released between March 2021 and June 2021 by the community project CAMEO (Continuous Automated Model Evaluation) [7]. These proteins contains less than 500 amino acid residues and have *<*25% sequence similarity with the SPOT-1D training set according to BlastClust.

**3.0.0.1 CASP** We used a collection of CASP [8] (Critical Assessment of protein Structure Prediction) datasets, especially the recent ones (e.g., CASP12 and CASP13). We used another dataset CASP-FM, comprising the CASP Free Modeling (FM) targets, which was previously compiled by the authors of SAINT. CASP-FM contains 56 domain sequences: 10 FM targets from CASP13, 22 FM targets from CASP12, 16 FM targets (out of 30) from CASP11 and 8 FM targets (out of 12) from CASP10.

### 4 Additional Results

**Figure S1:**
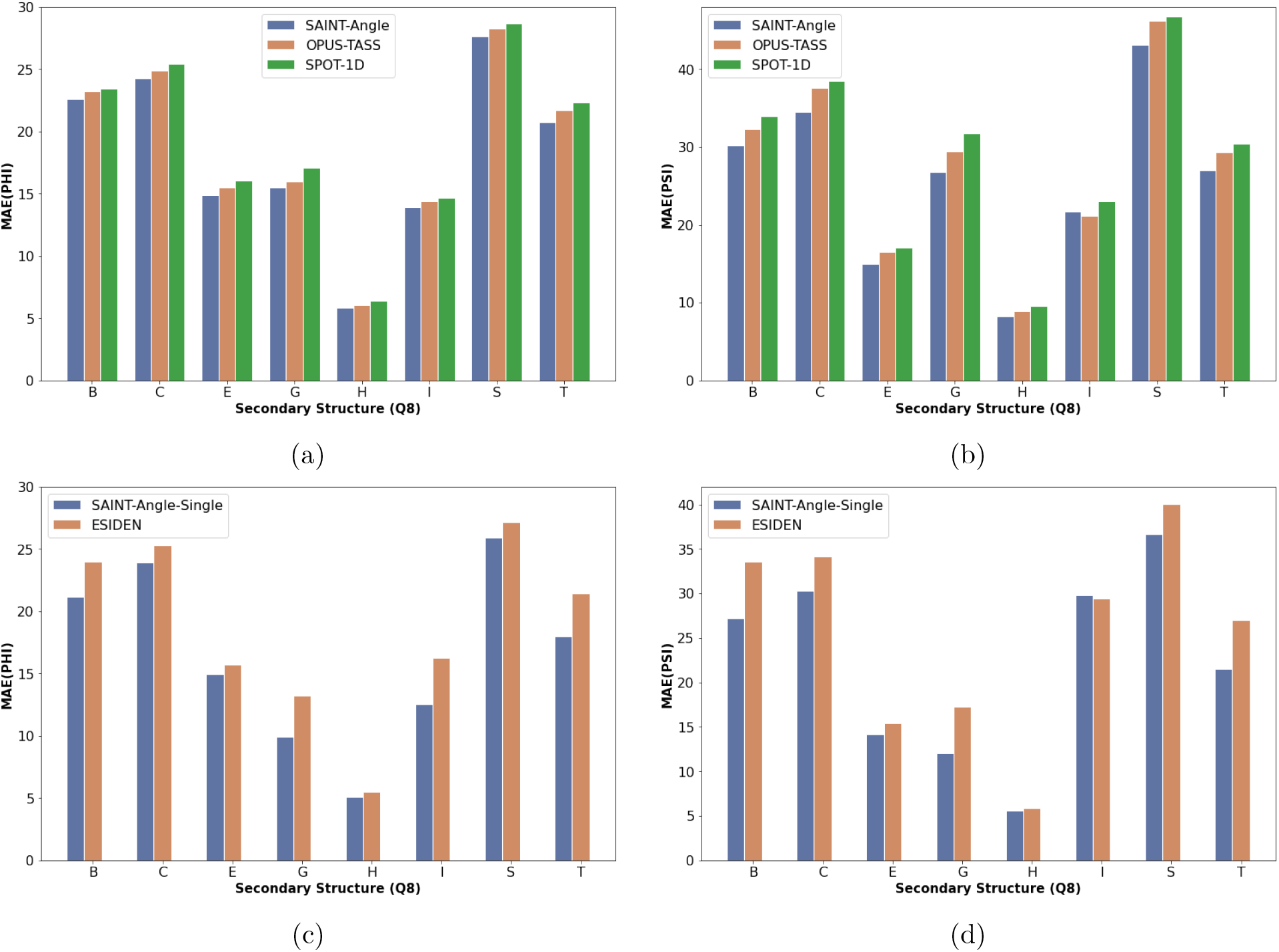
MAE(*ϕ*) and MAE(*ϕ*) of different methods on 8-class secondary structures for proteins in the TEST2016 dataset [5]. **(a)-(b)** MAE(*ϕ*) and MAE(*ψ*) of SAINT-Angle, OPUS-TASS, and SPOT-1D across different Q8 structures. **(c)-(d)** MAE(*ϕ*) and MAE(*ψ*) of SAINT-Angle-Single and ESIDEN across different Q8 structures.

## References

[1] Qian Jiang, Xin Jin, Shin-Jye Lee, and Shaowen Yao. Protein secondary structure prediction: A survey of the state of the art. Journal of Molecular Graphics and Modelling, 76:379–402, 2017.

[2] Mohammed AlQuraishi. End-to-end differentiable learning of protein structure. Cell systems, 8(4):292–301, 2019.

[3] Joe G Greener, Shaun M Kandathil, and David T Jones. Deep learning extends de novo protein modelling coverage of genomes using iteratively predicted structural constraints. Nature communications, 10(1):1–13, 2019.

[4] Jinbo Xu. Distance-based protein folding powered by deep learning. Proceedings of the National Academy of Sciences, 116(34):16856–16865, 2019.

[5] Andrew W Senior, Richard Evans, John Jumper, James Kirkpatrick, Laurent Sifre, Tim Green, Chongli Qin, Augustin Žídek, Alexander WR Nelson, Alex Bridgland, et al. Improved protein structure prediction using potentials from deep learning. Nature, 577(7792):706–710, 2020.

[6] Gang Xu, Qinghua Wang, and Jianpeng Ma. Opus-tass: a protein backbone torsion angles and secondary structure predictor based on ensemble neural networks. Bioinformatics, 36(20):5021–5026, 2020.

[7] GN Ramachandran, C Ramakrishnan, and V Sasisekharan. Stereochemistry of polypeptide chain configurations. Journal of Molecular Biology, 7(1):95–99, 1963.

[8] Aashish N Adhikari, Karl F Freed, and Tobin R Sosnick. De novo prediction of protein folding pathways and structure using the principle of sequential stabilization. Proceedings of the National Academy of Sciences, 109(43):17442–17447, 2012.

[9] Rhys Heffernan, Yuedong Yang, Kuldip Paliwal, and Yaoqi Zhou. Capturing non-local interactions by long short-term memory bidirectional recurrent neural networks for improving prediction of protein secondary structure, backbone angles, contact numbers and solvent accessibility. Bioinformatics, 33(18):2842–2849, 2017.

[10] Ofer Dor and Yaoqi Zhou. Real-spine: An integrated system of neural networks for real-value prediction of protein structural properties. PROTEINS: Structure, Function, and Bioinformatics, 68(1):76–81, 2007.

[11] Sitao Wu and Yang Zhang. Anglor: a composite machine-learning algorithm for protein backbone torsion angle prediction. PloS one, 3(10):e3400, 2008.

[12] Olav Zimmermann and Ulrich HE Hansmann. Support vector machines for prediction of dihedral angle regions. Bioinformatics, 22(24):3009–3015, 2006.

[13] Christopher Bystroff, Vesteinn Thorsson, and David Baker. Hmmstr: a hidden markov model for local sequence-structure correlations in proteins. Journal of molecular biology, 301(1):173–190, 2000.

[14] Rachel Karchin, Melissa Cline, Yael Mandel-Gutfreund, and Kevin Karplus. Hidden markov models that use predicted local structure for fold recognition: alphabets of backbone geometry. Proteins: Structure, Function, and Bioinformatics, 51(4):504–514, 2003.

[15] Rhys Heffernan, Kuldip Paliwal, James Lyons, Abdollah Dehzangi, Alok Sharma, Jihua Wang, Abdul Sattar, Yuedong Yang, and Yaoqi Zhou. Improving prediction of secondary structure, local backbone angles and solvent accessible surface area of proteins by iterative deep learning. Scientific reports, 5(1):1–11, 2015.

[16] Mike Schuster and Kuldip K Paliwal. Bidirectional recurrent neural networks. IEEE transactions on Signal Processing, 45(11):2673–2681, 1997.

[17] Chao Fang, Yi Shang, and Dong Xu. Prediction of protein backbone torsion angles using deep residual inception neural networks. IEEE/ACM transactions on computational biology and bioinformatics, 16(3):1020–1028, 2018.

[18] Christian Szegedy, Sergey Ioffe, Vincent Vanhoucke, and Alexander A Alemi. Inception-v4, inception-resnet and the impact of residual connections on learning. In Thirty-first AAAI conference on artificial intelligence, 2017.

[19] Michael Schantz Klausen, Martin Closter Jespersen, Henrik Nielsen, Kamilla Kjaergaard Jensen, Vanessa Isabell Jurtz, Casper Kaae Soenderby, Morten Otto Alexander Sommer, Ole Winther, Morten Nielsen, Bent Petersen, et al. Netsurfp-2.0: Improved prediction of protein structural features by integrated deep learning. Proteins: Structure, Function, and Bioinformatics, 87(6):520–527, 2019.

[20] Yujuan Gao, Sheng Wang, Minghua Deng, and Jinbo Xu. Raptorx-angle: real-value prediction of protein backbone dihedral angles through a hybrid method of clustering and deep learning. BMC bioinformatics, 19(4):73–84, 2018.

[21] Stephen F Altschul, Thomas L Madden, Alejandro A Schäffer, Jinghui Zhang, Zheng Zhang, Webb Miller, and David J Lipman. Gapped blast and psi-blast: a new generation of protein database search programs. Nucleic acids research, 25(17):3389–3402, 1997.

[22] Johannes Söding. Protein homology detection by hmm–hmm comparison. Bioinformatics, 21(7):951–960, 2005.

[23] Michael Remmert, Andreas Biegert, Andreas Hauser, and Johannes Söding. Hhblits: lightning-fast iterative protein sequence searching by hmm-hmm alignment. Nature methods, 9(2):173–175, 2012.

[24] Jack Hanson, Kuldip Paliwal, Thomas Litfin, Yuedong Yang, and Yaoqi Zhou. Improving prediction of protein secondary structure, backbone angles, solvent accessibility and contact numbers by using predicted contact maps and an ensemble of recurrent and residual convolutional neural networks. Bioinformatics, 35(14):2403–2410, 2019.

[25] Sepp Hochreiter and Jürgen Schmidhuber. Long short-term memory. Neural computation, 9(8):1735–1780, 1997.

[26] Kaiming He, Xiangyu Zhang, Shaoqing Ren, and Jian Sun. Deep residual learning for image recognition. In Proceedings of the IEEE conference on computer vision and pattern recognition, pages 770–778, 2016.

[27] Jack Hanson, Kuldip Paliwal, Thomas Litfin, Yuedong Yang, and Yaoqi Zhou. Accurate prediction of protein contact maps by coupling residual two-dimensional bidirectional long short-term memory with convolutional neural networks. Bioinformatics, 34(23):4039–4045, 2018.

[28] Yann LeCun, Léon Bottou, Yoshua Bengio, and Patrick Haffner. Gradient-based learning applied to document recognition. Proceedings of the IEEE, 86(11):2278–2324, 1998.

[29] Ashish Vaswani, Noam Shazeer, Niki Parmar, Jakob Uszkoreit, Llion Jones, Aidan N Gomez, Lukasz Kaiser, and Illia Polosukhin. Attention is all you need. In Advances in neural information processing systems, pages 5998–6008, 2017.

[30] Karim Lounici, Massimiliano Pontil, Alexandre B Tsybakov, and Sara Van De Geer. Taking advantage of sparsity in multi-task learning. arXiv preprint arXiv:0903.1468, 2009.

[31] Jens Meiler, Michael Müller, Anita Zeidler, and Felix Schmäschke. Generation and evaluation of dimension-reduced amino acid parameter representations by artificial neural networks. Molecular modeling annual, 7(9):360–369, 2001.

[32] Yong-Chang Xu, Tian-Jun ShangGuan, Xue-Ming Ding, and Ngaam J Cheung. Accurate prediction of protein torsion angles using evolutionary signatures and recurrent neural network. Scientific reports, 11(1):1–11, 2021.

[33] Mostofa Rafid Uddin, Sazan Mahbub, M Saifur Rahman, and Md Shamsuzzoha Bayzid. SAINT: self-attention augmented inception-inside-inception network improves protein secondary structure prediction. Bioinformatics, 36(17):4599–4608, 2020.

[34] Ahmed Elnaggar, Michael Heinzinger, Christian Dallago, Ghalia Rehawi, Yu Wang, Llion Jones, Tom Gibbs, Tamas Feher, Christoph Angerer, Martin Steinegger, et al. Prottrans: towards cracking the language of life’s code through self-supervised learning. bioRxiv, pages 2020–07, 2021.

[35] UniProt Consortium. The universal protein resource (uniprot). Nucleic acids research, 36(Suppl 1):D190–D195, 2007.

[36] Zihang Dai, Zhilin Yang, Yiming Yang, Jaime G Carbonell, Quoc Le, and Ruslan Salakhutdinov. Transformer-xl: Attentive language models beyond a fixed-length context. In Proceedings of the 57th Annual Meeting of the Association for Computational Linguistics, pages 2978–2988, 2019.

[37] Zhilin Yang, Zihang Dai, Yiming Yang, Jaime Carbonell, Russ R Salakhutdinov, and Quoc V Le. Xlnet: Generalized autoregressive pretraining for language understanding. In Advances in neural information processing systems, pages 5753– 5763, 2019.

[38] Jacob Devlin, Ming-Wei Chang, Kenton Lee, and Kristina Toutanova. BERT: Pre-training of deep bidirectional transformers for language understanding. In Proceedings of the 2019 Conference of the North American Chapter of the Association for Computational Linguistics: Human Language Technologies, Volume 1 (Long and Short Papers), pages 4171–4186, Minneapolis, Minnesota, June 2019. Association for Computational Linguistics.

[39] Chao Fang, Yi Shang, and Dong Xu. Mufold-ss: new deep inception-inside-inception networks for protein secondary structure prediction. Proteins: Structure, Function, and Bioinformatics, 86(5):592–598, 2018.

[40] Sergey Ioffe and Christian Szegedy. Batch normalization: Accelerating deep network training by reducing internal covariate shift. In International conference on machine learning, pages 448–456. PMLR, 2015.

[41] Y. Bengio, P. Simard, and P. Frasconi. Learning long-term dependencies with gradient descent is difficult. IEEE Transactions on Neural Networks, 5(2):157–166, 1994.

[42] Shuzhi Yu and Carlo Tomasi. Identity connections in residual nets improve noise stability. arXiv preprint arXiv:1905.10944, 2019.

[43] Diederik P Kingma and Jimmy Ba. Adam: A method for stochastic optimization. arXiv preprint arXiv:1412.6980, 2014.

[44] Guoli Wang and Roland L Dunbrack Jr. Pisces: a protein sequence culling server. Bioinformatics, 19(12):1589–1591, 2003.

[45] Frank Wilcoxon, SK Katti, and Roberta A Wilcox. Critical values and probability levels for the wilcoxon rank sum test and the wilcoxon signed rank test. Selected tables in mathematical statistics, 1:171–259, 1970.

[46] Sheng Wang, Jian Peng, Jianzhu Ma, and Jinbo Xu. Protein secondary structure prediction using deep convolutional neural fields. Scientific reports, 6(1):1–11, 2016.

[47] Jan Ludwiczak, Aleksander Winski, Antonio Marinho da Silva Neto, Krzysztof Szczepaniak, Vikram Alva, and Stanislaw Dunin-Horkawicz. Pipred–a deep-learning method for prediction of π-helices in protein sequences. Scientific reports, 9(1):1– 9, 2019.

## References

[1] Jack Hanson, Kuldip Paliwal, Thomas Litfin, Yuedong Yang, and Yaoqi Zhou. Improving prediction of protein secondary structure, backbone angles, solvent accessibility and contact numbers by using predicted contact maps and an ensemble of recurrent and residual convolutional neural networks. Bioinformatics, 35(14):2403–2410, 2019.

[2] Mostofa Rafid Uddin, Sazan Mahbub, M Saifur Rahman, and Md Shamsuzzoha Bayzid. SAINT: self-attention augmented inception-inside-inception network improves protein secondary structure prediction. Bioinformatics, 36(17):4599–4608, 2020.

[3] Gang Xu, Qinghua Wang, and Jianpeng Ma. Opus-tass: a protein backbone torsion angles and secondary structure predictor based on ensemble neural networks. Bioinformatics, 36(20):5021–5026, 2020.

[4] Yong-Chang Xu, Tian-Jun ShangGuan, Xue-Ming Ding, and Ngaam J Cheung. Accurate prediction of protein torsion angles using evolutionary signatures and recurrent neural network. Scientific reports, 11(1):1–11, 2021.

[5] Jack Hanson, Kuldip Paliwal, Thomas Litfin, Yuedong Yang, and Yaoqi Zhou. Accurate prediction of protein contact maps by coupling residual two-dimensional bidirectional long short-term memory with convolutional neural networks. Bioinformatics, 34(23):4039–4045, 2018.

[6] Helen M Berman, John Westbrook, Zukang Feng, Gary Gilliland, Talapady N Bhat, Helge Weissig, Ilya N Shindyalov, and Philip E Bourne. The protein data bank. Nucleic acids research, 28(1):235–242, 2000.

[7] Juergen Haas, Steven Roth, Konstantin Arnold, Florian Kiefer, Tobias Schmidt, Lorenza Bordoli, and Torsten Schwede. The protein model portal—a comprehensive resource for protein structure and model information. Database, 2013, 2013.

[8] Patrice Koehl and Michael Levitt. A brighter future for protein structure prediction. nature structural biology, 6(2):108–111, 1999.

